# Identification of Family-Specific Features in Cas9 and Cas12 Proteins: A Machine Learning Approach Using Complete Protein Feature Spectrum

**DOI:** 10.1101/2024.01.22.576286

**Authors:** Sita Sirisha Madugula, Pranav Pujar, Nammi Bharani, Shouyi Wang, Vindi M. Jayasinghe-Arachchige, Tyler Pham, Dominic Mashburn, Maria Artilis, Jin Liu

## Abstract

The recent development of CRISPR-Cas technology holds promise to correct gene-level defects for genetic diseases. The key element of the CRISPR-Cas system is the Cas protein, a nuclease that can edit the gene of interest assisted by guide RNA. However, these Cas proteins suffer from inherent limitations like large size, low cleavage efficiency, and off-target effects, hindering their widespread application as a gene editing tool. Therefore, there is a need to identify novel Cas proteins with improved editing properties, for which it is necessary to understand the underlying features governing the Cas families. In the current study, we aim to elucidate the unique protein attributes associated with Cas9 and Cas12 families and identify the features that distinguish each family from the other. Here, we built Random Forest (RF) binary classifiers to distinguish Cas12 and Cas9 proteins from non-Cas proteins, respectively, using the complete protein feature spectrum (13,495 features) encoding various physiochemical, topological, constitutional, and coevolutionary information of Cas proteins. Furthermore, we built multiclass RF classifiers differentiating Cas9, Cas12, and Non-Cas proteins. All the models were evaluated rigorously on the test and independent datasets. The Cas12 and Cas9 binary models achieved a high overall accuracy of 95% and 97% on their respective independent datasets, while the multiclass classifier achieved a high F1 score of 0.97. We observed that Quasi-sequence-order descriptors like Schneider-lag descriptors and Composition descriptors like charge, volume, and polarizability are essential for the Cas12 family. More interestingly, we discovered that Amino Acid Composition descriptors, especially the Tripeptide Composition (TPC) descriptors, are important for the Cas9 family. Four of the identified important descriptors of Cas9 classification are tripeptides PWN, PYY, HHA, and DHI, which are seen to be conserved across all the Cas9 proteins and were located within different catalytically important domains of the Cas9 protein structure. Among these four tripeptides, tripeptides DHI and HHA are well-known to be involved in the DNA cleavage activity of the Cas9 protein. We therefore propose the the other two tripeptides, PWN and PYY, may also be essential for the Cas9 family. Our identified important descriptors enhanced the understanding of the catalytic mechanisms of Cas9 and Cas12 proteins and provide valuable insights into design of novel Cas systems to achieve enhanced gene-editing properties.

## 1. Introduction

Clustered Regularly Interspaced Short Palindromic Repeats(CRISPR) and its associated proteins (Cas) together form the CRISPR-Cas system that functions as part of an adaptive prokaryotic immune system^1–7^. Its discovery and successive development into gene-editing technology hold a promise to correct gene-level defects for several genetic diseases. The Cas proteins, together with guide RNA (gRNA), form a programmable system to recognize virtually any target DNA sequence preceding a signature Protospacer Adjacent Motif [PAM, a short DNA motif that is recognized sequence-specifically by Cas gRNA to initiate binding to DNA]^8,9^ Due to its ability to edit any gene and its straightforward design, this technology has garnered immense attention as potential therapeutic to address various genetic disorders like Haemophilia, beta-thalassemia, Amyotrophic Lateral Sclerosis, Muscular Dystrophy, and different types of cancer^10–12^. Recently, U.S. Food and Drug Administration (FDA) approved “exa-cel”, the first CRISPR treatment for sickle cell disease, by Vertex and CRISPR Therapeutics soon after the U.K. approved exa-cel for sickle cell disease in November 2023^13^. Despite these advancements in CRISPR/Cas technology, limitations like low specificity, off-target effects, bigger size of Cas and inefficient delivery of CRISPR-Cas components remain significant challenges for Cas proteins impeding their widespread application. Therefore, it is essential to discover novel Cas proteins with better editing capabilities and a reduced size for efficient delivery, aiming to develop CRISPR-Cas systems with higher accuracy in gene editing. To achieve this, an understanding of the underlying characteristics of each Cas subfamily is essential.

Different Cas proteins have different functions and participate in different stages of the CRISPR-Cas system^14^. According to the number of Cas proteins involved in effector complex formation, CRISPR-Cas systems are divided into two main classes. While class 1 (I, III, and IV) utilize multiple Cas proteins to generate the effector complex for recognition and cleavage, Class 2 (II, V, and VI) utilize a single Cas protein^15–20^. Compared to Class 1, the simpler architecture of effector complexes in Class 2 CRISPR-Cas systems offers more advantages for innovative gene editing technologies^21,22^. Some common Class 2 effectors include Cas9, Cas12a, Cas12b, Cas13a, Cas13b and Cas13c^5,23^.

Presently, the existing protein database like UniProt^24^ has limited annotated Cas structures with comprehensive information about their structure, function, and characteristic domains. With respect to the CRISPR/Cas systems, CRISPRFinder^25^, PILER-CR^26^, CRISPRDetect^27^, CRISPRcasIdentifier^28^ represent some of the available computational and Artificial Intelligence (AI) based tools. While these existing computational tools predominantly operate as web-based packages, focusing on the identification of CRISPR arrays, there is a scarcity of computational tools specifically designed to identify and characterize novel Cas proteins. In this regard, HMMCAS^29^; a Hidden Markov Model (HMM) based tool is one of the earliest tool developed by Chai et.al in 2019 for Cas functional domain characterization and identification. Subsequently, Yang and colleagues developed CASPredict^30^, a web-based tool built on Support Vector Machine (SVM) using Optimal DPC (ODPC) descriptors to identify new Cas proteins, achieving an overall accuracy of 84%. Notably, the tool was designed with a focus on optimal dipeptide descriptors, potentially leaving room for improvement by incorporating additional descriptors encoding coevolutionary, structural, positional, and other sequential information to comprehensively represent Cas proteins. Enhancements in these aspects could capture new features that can guide in designing novel Cas proteins with optimized size, enhanced specificity, and improved cleavage properties.

In the field of CRISPR-Cas systems, Cas9 has gained prominence due to its swift and cost-effective cleavage activity, positioning it favorably for applications in diverse fields beyond its traditional use.^31^ Meanwhile, the Cas12 family, with its unique collateral cleavage properties and expanded capabilities, broadens the toolkit for precise genetic editing, offering enhanced versatility in molecular applications^32–34^. In the current study, we aim to identify the characteristic protein descriptors specific to Cas9 and Cas12 families and delineate the distinct features that define each family. For this purpose, we developed Machine Learning (ML) based pipeline (Figure 1) to classify Cas12 and Cas9 protein families. The pipeline includes Random Forest (RF) classification models built using the complete spectrum of protein features encompassing 13,495 descriptors which account for the various structural, functional, physiochemical, and coevolutionary information of Cas12 and Cas9 proteins. We developed two binary classifiers for Cas9 vs. Non-Cas and Cas12 vs. Non-Cas classifications and a multiclass classifier to differentiate between Cas9, Cas12 and Non-Cas proteins (represented as “Cas9-Cas12-Non-Cas” hereafter). In this process, we also identified the specific descriptors unique to each protein family. Our study represents a pioneering effort in dedicated Cas9 and Cas12 classification, utilizing an extensive array of protein descriptors. While there are existing classification tools, our approach stands out by comprehensively delineating Cas family-specific features. These insights not only contribute to the existing body of knowledge but also provide a roadmap for designing improved Cas proteins with optimized size, enhanced cleavage efficiency, and reduced off-target effects; an endeavor crucial for advancing gene-editing applications.

**Figure 1.**
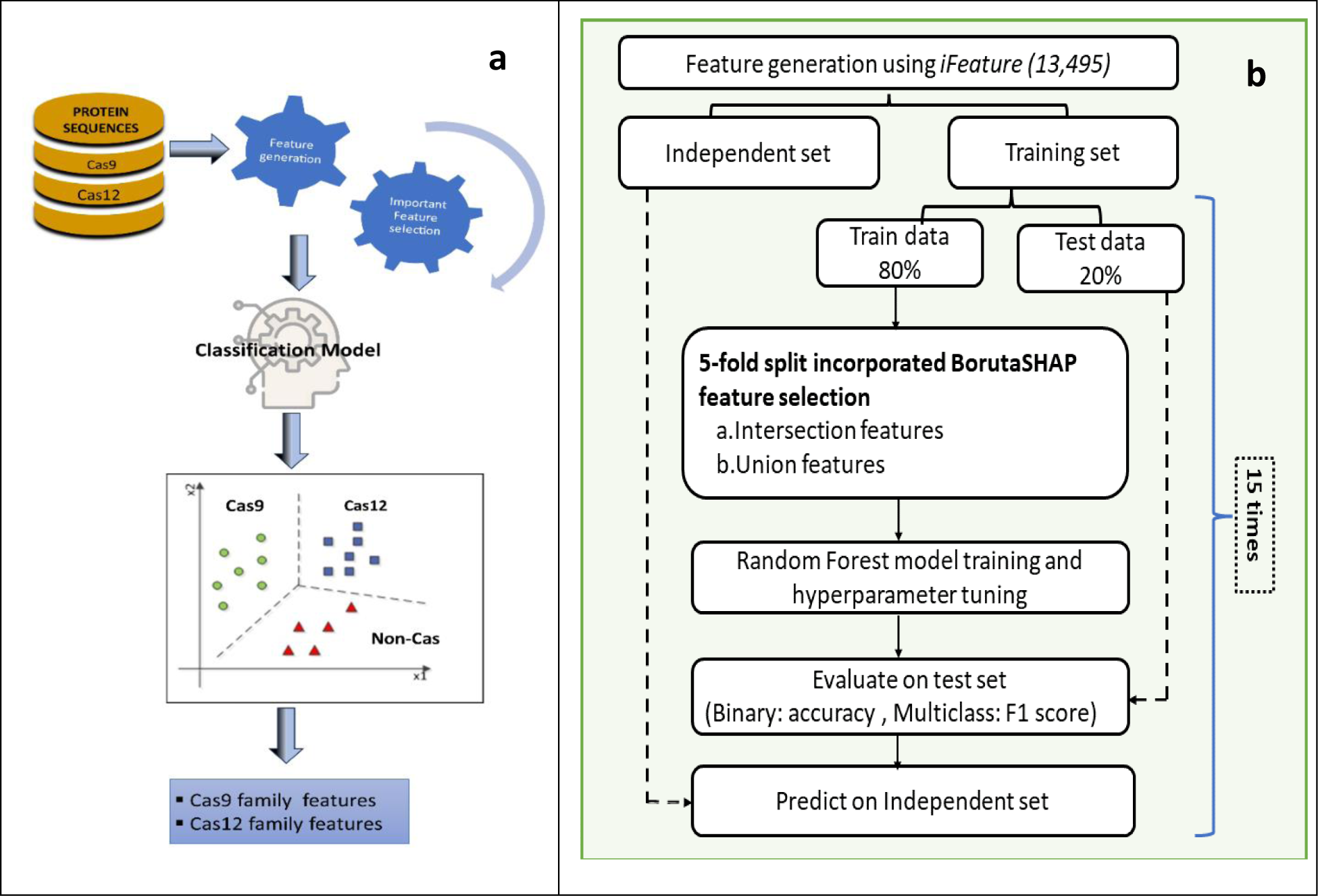
Workflow of the study. **(a)** Overview of the study depicting the important steps of feature generation, feature selection followed by classification. The important features characteristic of the Cas9 and Cas12 families are further analyzed. **(b)** The detailed steps involved of our ML pipeline. Protein features calculated in the feature generation step are subjected to a 5-fold BorutaSHAP feature selection method to identify the important features. Intersection of accepted features across all 5 folds is taken to get the intersection features. Similar procedure is repeated across All 5 folds by taking union to obtain union features. The important intersection features are further used to develop RF classifiers which are then used to make predictions on the independent set.

## 2. Materials and Methods

### 2.1. Data sources

#### a) Cas12 dataset

##### Training data

The Cas12 training set contains 288 proteins which was prepared using the Cas12 proteins collected from different sources including UniProt^23^, RCSB-PDB^35^ and literature which were manually inspected for the presence of the characteristic Cas12 protein domains.

##### Independent data

Independent dataset includes 41 Cas proteins are mutated forms and engineered versions of original Cas12 proteins which were obtained from the literature.

#### b) Cas 9 dataset

##### Training data

The Cas9 training dataset contains 837 Cas proteins which were collected from InterPro database^36^ from annotated protein entries and were manually inspected for the presence of domains characteristic to Cas9.

##### Independent data

This set contains 40 Cas proteins which are engineered Cas9 collected from literature.

#### c) Non-Cas dataset

##### Training data

The dataset contains 600 Non-Cas proteins which included proteases and exonucleases.

##### Independent data

The non-Cas constituted of proteins collected from diverse families viz., DNA endonucleases, proteases, exonucleases, and helicases. We also included other types of Cas proteins such as Cas2, Cas5, Cas8 and Cas13 to develop robust models capable of distinguishing Cas12 and Cas9 proteins from these proteins. The three datasets used in the study are summarized in Table 1.

**Table 1.** Summary of the number of data points in different datasets used in the study.

| Dataset | Training | Independent |
| --- | --- | --- |
| Cas12 | 288 | 41 |
| Cas 9 | 837 | 40 |
| Non-Cas | 600 | 48 |
| Total | **1725** | **129** |

### 2.2. Feature generation

To calculate the protein descriptors(or features), we employed iFeature^37^ an open-source Python package, that can generate a plethora of protein and peptide descriptors from their amino acid sequences. The generated descriptors belong to various categories viz. sequential, physicochemical, structural, and evolutionary descriptors representing the complete profile of a protein. For this study, we calculated descriptors belonging to seven groups (Table 2) from the range of descriptor categories offered by the program. These seven groups are the only descriptors that can be computed using FASTA sequences of varying lengths. Every descriptor group consists of multiple descriptor sets, and each descriptor set in turn contains several descriptors. The seven groups comprise a total of 13,494 descriptors which are calculated for Cas9 and Cas12 datasets.

**Table 2.** List of the descriptors used in the study.

| Descriptor Group | Descriptor set | No. of Descriptors |
| --- | --- | --- |
| Amino Acid Composition | Amino acid composition (AAC) | 20 |
|  | Composition of k-spaced amino acid pairs (CKSAAP) | 2400 |
|  | Dipeptide composition (DPC) | 400 |
|  | Dipeptide deviation from expected mean (DDE) | 400 |
|  | Tripeptide composition (TPC) | 8000 |
| Grouped Amino Acid Composition | Grouped amino acid composition (GAAC) | 5 |
|  | Composition of k-spaced amino acid group pairs (CKSAAGP) | 150 |
|  | Grouped dipeptide composition (GDPC) | 25 |
|  | Grouped tripeptide composition (GTPC) | 125 |
| Autocorrelation | Moran (Moran) | 240 |
|  | Geary (Geary) | 240 |
|  | Normalized Moreau-Broto (NMBroto) | 240 |
| C/T/D | Composition (CTDC) | 39 |
|  | Transition (CTDT) | 39 |
|  | Distribution (CTDD) | 195 |
| Conjoint Triad | Conjoint triad (CTriad) | 343 |
|  | Conjoint *k*-spaced triad (KSCTriad) | 343 |
| Quasi-Sequence-Order(QSO) | Sequence-order-coupling number (SOCNumber) | 60 |
|  | Quasi-sequence-order descriptors (QSOrder) | 100 |
| Pseudo-Amino-Acid Composition | Pseudo-amino-acid composition (PAAC) | 50 |
|  | Amphiphilic PAAC (APAAC) | 80 |

Table adopted from iFeature package.

### 2.3. Feature selection

Since a large feature set is considered in this study, feature selection algorithms must be applied to avoid multicollinearity and identify contributing features. For this purpose we applied BorutaSHAP^38^ feature selection algorithm which can handle multicollinearity among features by applying SHapley Additive exPlanations (SHAP) analysis, and utilizes a game theoretic approach to compute the contribution of each feature towards the prediction^39,40^. To identify consistently contributing features and avoid bias, we applied a stratified 5-fold split of the training data used in BorutSHAP algorithm. This generated 5 sets of accepted features for the 5 folds at each run. At this stage, an intersection is taken over the accepted features across all the 5 folds creating the intersection feature set. Likewise, a union of all the accepted features across the 5 folds is taken to construct the union feature set. RF models were constructed using both sets of features separately. However, we noticed that the performance of models trained on intersection features is almost comparable to the performance of models trained using union features. Therefore, we discuss the intersection models’ performance in the forthcoming sections as they explain the model using fewer features compared to union.

### 2.4. Evaluation metrices

The performance of our models is evaluated using six different evaluations metrices; accuracy, recall, F1 score, Precision, specificity and Area Under The Receiver Operating Characteristic Curve (AUC-ROC) on the test dataset. For the multiclass classification of Cas9-Cas12-Non-Cas proteins, F1 score is chosen to be a suitable metric to evaluate the performance. When dealing with imbalanced datasets such as the data used in this study, overall accuracy is considered misleading due to the influence of majority class on the training procedure leading to improper learning of the minority class. On the other end, F1 score which combine precision and recall are thought to provide better assessment since it considers both the rate of correctly classified positives (True Positives) and the falsely classified positives (False Positives).

### 2.5. Experimental Settings

Three different classification tasks were carried out in this study. These include Cas12 vs. Non-Cas and Cas9 vs. Non-Cas binary classifications and a multiclass classifications of Cas12 vs.Cas9 vs. Non-Cas proteins. RF was employed for model building in all classifications. The training data is split using “stratified” data split in 80-20 ratio to generate training and test datasets respectively. The training dataset was next used to carry out feature selection using BorutaSHAP algorithm to identify the important descriptors. Scikit-Learn’s GridSearchCV^41^ with a 5-fold CV is used for hyperparameter tuning in which a grid of different parameter ranges is specified and an exhaustive search is done over all the specified values to identify the combination of parameters producing best fit. In all the three classifications, we tuned 2 parameters namely “n_estimators” and “max_depth” which are the number of trees in the forest and the maximum depth of the trees respectively. **Table S1** describes the hyperparameters and their ranges used to tune the RF models. Accuracy was used to evaluate the performance of cross-validated models during training for binary classifiers while F1 score was used for the multiclass classification.

The class imbalance in the training dataset was handled by giving “class weights=balanced” keyword which assigns weights to the individual classes such that the weights are inversely proportional to their class frequency. This procedure avoids bias towards the majority class ensuring that the model learns equal representation of all classes. To observe the variance of our models we repeated the steps of feature selection, model training, evaluation, and test set prediction over 15 different data splits. For further evaluation, the developed models were also predicted on their respective independent datasets and an average of the evaluation matrices over the 15 splits is reported.

## 3. Results

### 3.1. Cas12 and Cas9 Binary Classification

To understand the molecular features governing the Cas12 and Cas9 families, we developed binary classifiers that can identify the two types of Cas proteins. Since we considered large number of descriptors, we used the Random Forest (RF) algorithm to handle multicollinearity. The RF algorithm utilizes many decision trees trained with either bagging or boosting methods, yielding an outcome based on the average of predictions from all trees. More trees lead to more robust models. By using a minimum number of descriptors, the algorithm minimizes overfitting and improves precision. Respective RF binary models were built on Cas12 and Cas9 training datasets and extensively evaluated on the respective test sets followed by independent datasets.

The feature selection to evaluation experiments were repeated 15 times with different random data splits to record the mean performance and standard deviation of our models built using intersection features across the 5-fold cross validation in each trial. The experimental results are summarized in Tables 3 and 4, respectively.

**Table 3:** Test set performance of the three classification models using intersection features averaged over 15 runs.

| Classification type | Accuracy | Precision | Recall | F1-score | AUC | Specificity |
| --- | --- | --- | --- | --- | --- | --- |
| Cas12 vs. Non-Cas | 0.9636±<br>0.014 | 0.9634±<br>0.016 | 0.9537±<br>0.019 | 0.9581±<br>0.017 | 0.9537±<br>0.019 | 0.9822±<br>0.011 |
| Cas9 vs. Non-Cas | 0.9969±<br>0.003 | 0.9972±<br>0.002 | 0.9965±<br>0.003 | 0.9968±<br>0.003 | 0.9965±<br>0.003 | 0.9938±<br>0.007 |
| Cas12-Cas9-Non-Cas | 0.9980±<br>0.002 | 0.9984±<br>0.001 | 0.9984±<br>0.002 | 0.9984±<br>0.002 | 0.9996±<br>0.0008 | 0.9988±<br>0.001 |

**Table 4:** Independent set performance of the three classification models using intersection features averaged over 15 runs.

| Classification type | Accuracy | Precision | Recall | F1-score | AUC | Specificity |
| --- | --- | --- | --- | --- | --- | --- |
| Cas12 vs. Non-Cas | 0.9222±<br>0.050 | 0.9392±<br>0.038 | 0.9159±<br>0.055 | 0.9197±<br>0.054 | 0.915±<br>0.055 | 0.9945±<br>0.011 |
| Cas9 vs. Non-Cas | 0.9564±<br>0.030 | 0.9567±<br>0.028 | 0.9554±<br>0.030 | 0.9554±<br>0.031 | 0.955±<br>0.033 | 0.9625±<br>0.011 |
| Cas12-Cas9-Non-Cas | 0.9794±<br>0.008 | 0.9795±<br>0.007 | 0.9801±<br>0.009 | 0.9796±<br>0.008 | 0.996±<br>0.001 | 0.9896±<br>0.004 |

As seen in Tables 3 and 4, all the three models performed well across both the test and independent datasets. Since Cas12 vs. Non-Cas and Cas9 vs. Non-Cas are binary classification problem, accuracy is considered as a suitable metric for model evaluation. However, we also calculated other metrices like precision, recall and F1 scores for a more accurate representation of the model performance.

On test sets, Cas12 and Cas9 binary models achieved a high average accuracy of over 96% and 99% for respectively. Along with the accuracy, the models also achieved high values of precision, recall and F1 scores as shown in Tables 3. On the independent dataset (Table 4), the Cas12 models achieved a high average accuracy of close to 97% along with a high average precision, specificity and F1 scores of over 96%. Likewise, the Cas9 models also performed well on the independent datasets and achieved high average precision, recall and F1 scores above 97%. Therefore, our Cas12 and Cas9 classification models achieved a high performance on both test and independent datasets.

### 3.2. Cas12-Cas9-Non-Cas Multiclass Classification

In this study, we developed a multiclass classification model that can differentiate between Cas12, Cas9 and Non-Cas proteins. The multiclass models can be useful to identify and validate if a newly designed protein is Cas9, Cas12 or a Non-Cas protein. As seen in Table 3 and 4, the models achieved an extremely high average performance around 99% across all the metrices over the 15 splits on the test set. On the other hand, the multiclass models also performed extremely well on the independent datasets showing a high average F1 score of about 98% along with a high average recall and AUC.

For each of the three classification models, we utilized the best performing model among the 15 splits to discuss the classification performance on their respective independent datasets in all the forthcoming sections. On the Cas12 independent dataset, the best Cas12 RF model correctly classified all Non-Cas proteins and correctly categorized 39 out of 41 Cas proteins, resulting in the higher precision and slightly lower recall. Alternatively, on the Cas9 independent dataset, the best Cas9 RF model classified 39 out of 40 as Cas9 thereby misclassifying only one protein. The model also classified 46 out of 48 non-Cas proteins correctly, producing only 2 false positives. In the multiclass model all Cas12 proteins were correctly classified while missing only one for Cas9 and two for Non-Cas proteins. The models misclassified only one Cas9 as Non-Cas (FN) and two Non-Cas proteins as Cas9 (FP) which shows the model’s low error rate and efficient performance on the independent set. These results demonstrate the high performance of our models developed using the complete spectrum of protein features.

### 3.3. Description of the important features of the two Cas families

To identify descriptors that characterize each Cas family, a detailed analysis of the intersection features obtained across the 15 splits is performed and the top-ranking descriptors are identified in each classification task. All descriptors are assigned ranks based upon their “average feature importance” obtained from BorutaSHAP feature selection algorithm. A global rank is then calculated for every descriptor by averaging over its rank across all 15 splits. This global rank is then used to identify the top-ranking descriptors.

#### 3.3.1. Important Features to Discriminate Cas12 vs. Non-Cas

BorutaSHAP method identified 93 important descriptors for Cas12 models (Figure 2a). The 93 descriptors belong to different descriptor sets of Table 3. From Figure 2a, maximum number of descriptors belong to the SOCNumber descriptor set followed by other descriptor sets. SOCNumber denoting the sequence order information of proteins often exhibits correlations with their biological functions. Consequently, it finds applications in tasks such as protein subcellular location identification^42^. Composition (C), Transition (T), and Distribution (D) (C/T/D) descriptors elucidate information about the prevalence, changes, and overall arrangement of amino acid patterns with specific structural or physicochemical properties within protein sequences. They are instrumental in elucidating folding patterns in proteins^43^, contributing significantly to the understanding of protein structure and function. CKSAAGP and GDPC are a subset of the Grouped Amino Acid Composition (GAAC) descriptor set that describes the frequency of amino acid groups derived based on shared physiochemical properties which are again directly related to the biological functions of the protein sequences. Similarly, AAC descriptors also describe the frequency of individual amino acids in the protein sequences. Therefore, these categories are commonly used for various protein or peptide prediction related tasks.^44–48^

**Figure 2.**
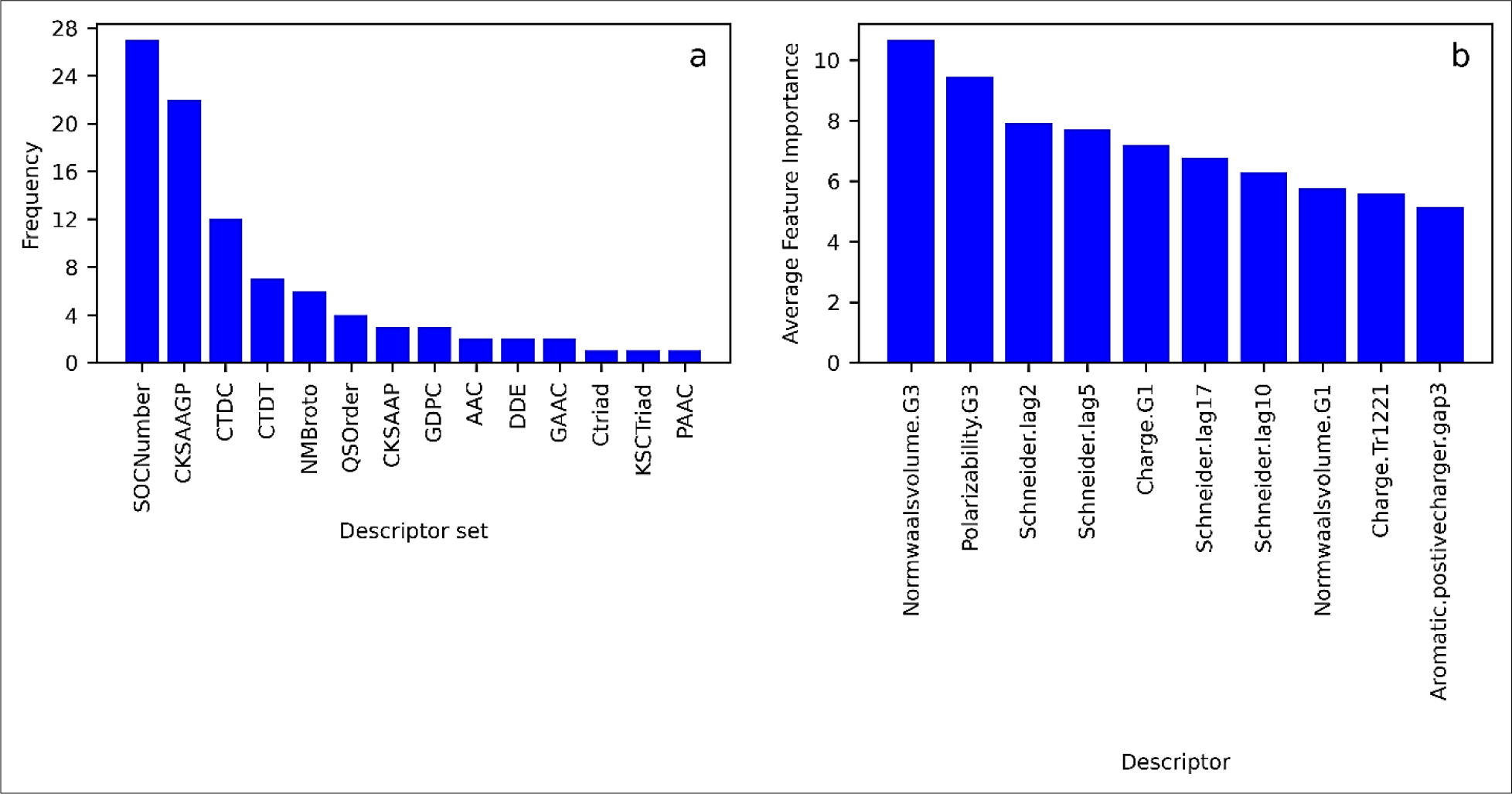
Summary of the important features in CAS12 vs. Non-CAS classification. **(a)** Descriptor set wise distribution of important features identified in Cas12 vs. Non-Cas RF model. X axis represents the descriptor groups to which the important features belong to and y-axis represents a count of the important descriptors within the said descriptor groups **(b)** Top 10 descriptors of Cas12 models.

##### Analysis of Cas12 top 10 ranking important descriptors

To gain further insights into the top features governing Cas12 family, we further analyzed the top 10 important descriptors (Figure 2b). The detailed analysis is summarized as follows:

- The top 10 descriptors with highest feature importance are mostly (C/T/D), Quasi-Sequence-Order (QSO), and Grouped Amino Acid Composition (GAAC) category. The latter two groups represent the significance of global content of different amino acid groups and the order of amino acid residues within Cas12 sequences respectively.
- The C/T/D descriptors represent the Composition, Transition, and Distribution patterns of a specific physicochemical property in protein sequences^44–46^. Accordingly, the 20 amino acids are categorized into three main groups based on seven physicochemical attributes namely hydrophobicity, Van der Waals Volume, polarity, polarizability, charge, secondary structures, and solvent accessibility together accounting for total C/T/D descriptors^47,48^.Composition descriptors (CTDC), represent the global composition of the three encoded amino acid groups of a particular property within the complete protein sequence.
- The top contributing descriptor “normwaalsvolume.G3” in Figure 2b is a CTDC descriptor describing the importance of group-3 normalized Van der Waals Volume residues (M,H,K,F,R,Y,W) having large residual volume in the Cas12 protein sequences. These bulky residues in protein sequences are usually associated with bulkiness of the protein structure which may have implications on the three-dimensional structure and functional sites of the protein. In the Cas12 sequences, this observation signifies that these residues contribute to the overall bulkiness of Cas12 protein. Further investigations would be needed to precisely interpret the functional significance of this descriptor in Cas12 proteins.
- Similarly, the second most important descriptor ‘Polarizability.G3’ (Figure 2b) is also a CTDC descriptor that describes the global distribution pattern of group-3 polarizability residues within the Cas12 sequences. C/T/D descriptors were first developed b*y* Cai et al.^54–55^. “Polarizability.G3” feature describes the importance of the global composition of the group-3 polarizability residues (K, M, H, F, R, Y, W) within the Cas12 family. All these seven residues also have large polarizability values between 0.219 and 0.409. This feature correlates ell with large polarizability of active site residues in proteins being known to facilitate easier protein-ligand interactions, which is a vital requirement for the biological functioning of proteins^51–53^. Interestingly, the mechanism of dsDNA cleavage by Cas12 proteins involves the action of three conserved lysine residues (a group-3 polarizability residue) whose positioning is critical for the unwinding of helical dsDNA^54^. Therefore, this descriptor is important for the Cas12 proteins and could be insightful in their de-novo design activities.
- Along with polarizability and normalized Van der Waals Volume, we also observed that charge-based CTDC descriptors including charge.G1 and Charge.Tr1221 are within the top descriptors, suggesting the importance of the physiochemical properties such as charge in the Cas12 proteins.
- We also observed a significant number of “Schneider lag” descriptors (lag2, lag5, lag10, and lag17) among the top 10 descriptors for the Cas12 family. These descriptors belong to the SOCNumber set, which represents the correlation modes between amino acids. It shows the associations between amino acids in different regions along the protein sequence, as depicted by their sequence-order-coupling (SOC) numbers between pairs of residues separated by a distance. SOC numbers give information on the coupling mode between contiguous residues separated by lag distance along the protein sequence. The *d*^th^ rank SOC number between pair of residues is given below:

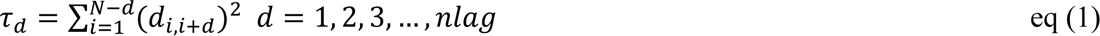

Where d_i_, d_i+d_ are distance entries in Schneider and Grantham physiochemical distance matrices indicating the distance between i^th^ and i+d^th^ residues which are derived based upon the residue physiochemical properties including hydrophobicity, hydrophilicity, polarity, and side-chain volume. Nlag denotes the maximum value of the lag (default value: 30) and N is the length of a protein or peptide sequence. SOCNumber descriptors were first introduced by Schneider and Wrede^55^ and Grantham who derived distance metrices between pairs of the 20 amino acids based upon their physiochemical properties described above. Since the distance from residue i to residue i+d is generally not equal to that from i+d to i; descriptors derived from such an approach can effectively distinguish the directionality of protein sequence order. Kuo-Chen Chou then used the former distance matrix to incorporate sequence order effect and predicted subcellular location of the proteins^42,56^. Therefore, sequence-order-information can be employed to predict different protein attributes. Larger lag-number descriptors indicate associations between residues lying far apart along the protein sequence length which could be interpreted as long-range contacts within the residues in their 3D protein structure or may also be associated with certain protein-protein interactions. This descriptor is also related to the co-evolutionary information of amino acids in protein sequences which influences its protein folding and 3D structure which in turn determines its biological function. The descriptor finds relevance in Cas12 vs. Non-Cas binary classification by providing insights into the evolutionary trajectory of Cas proteins.

#### 3.3.2. Important Features to Discriminate CAS9 vs. Non-CAS

For the Cas9 family, BorutaSPHAP gave 108 important descriptors seen in Figure 3a. Unlike Cas12, DDE, which are amino acid composition type descriptors, emerged as the descriptor set with the highest number of important features in Cas9 family. This is followed by SOCNumber, TPC, and other sets shown in Figure 3a. While there are similarities between the important descriptor sets of both Cas families, some sets like Distribution (CTDD), Pseudo amino acid composition (APAAC), and Autocorrelation descriptors like Geary and Moran type of descriptors are seen only in the Cas9 family. Autocorrelation descriptors define the correlation between two proteins in terms of specific physiochemical properties derived from the distribution patterns of amino acid properties along the protein sequences. These descriptors can quantify patterns in physicochemical properties along protein sequences which can be correlated to biological function. Autocorrelation descriptors are employed in a wide range of applications involving predicting various protein-protein interactions and identifying gene-functions.^57–59^ It should be noted that although DDE set had the highest count of important descriptors given by BorutaSHAP, it is the TPC descriptors that hold higher values of average feature importance in the Cas9 vs. Non-Cas classification making them majority in the top 10 descriptors (Figure 3b)

**Figure 3.**
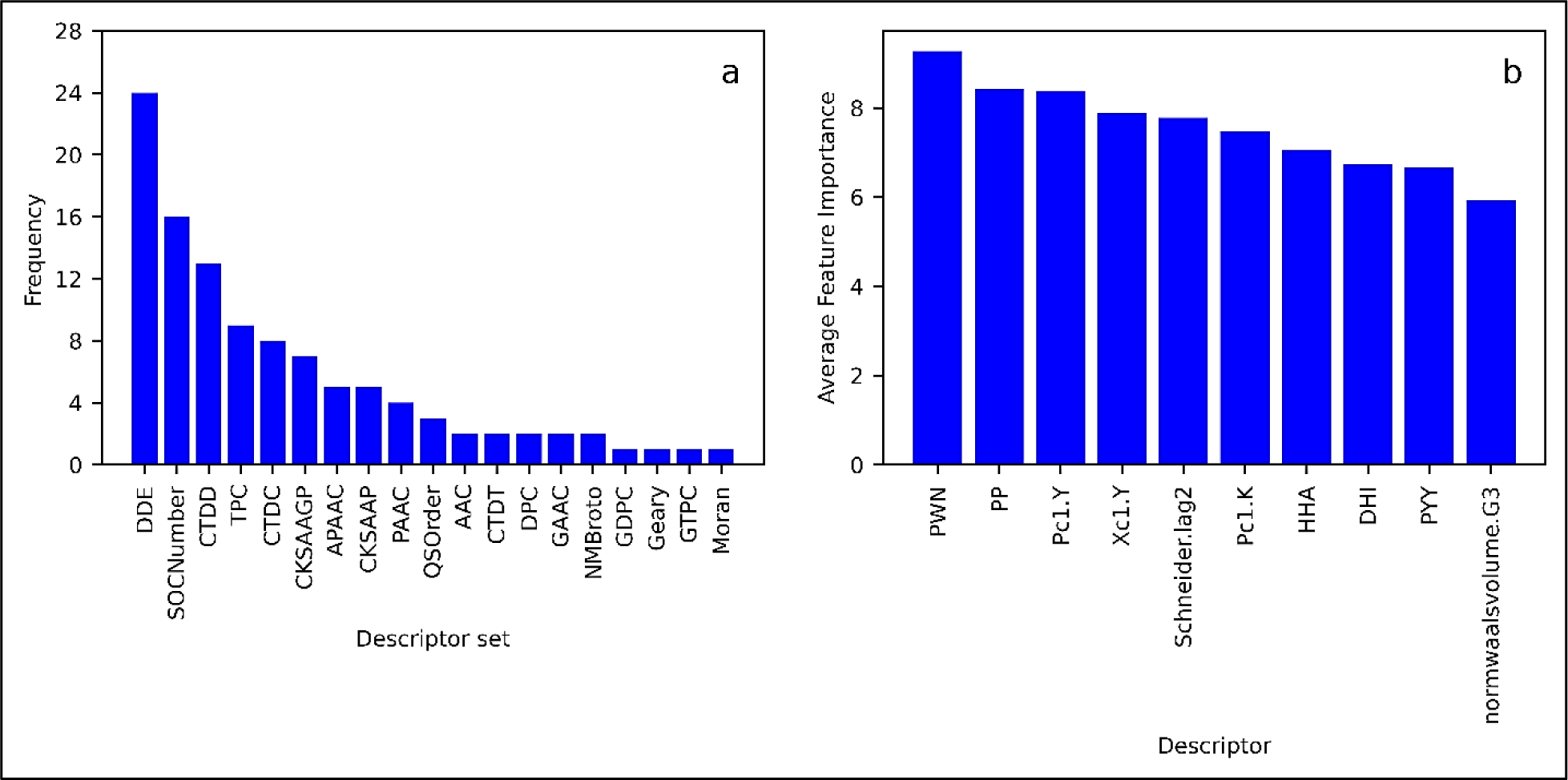
Summary of the important features in CAS9 vs. Non-CAS classification **(a)** Descriptor set wise distribution of important features identified in Cas9 RF model. X-axis represents the descriptor groups to which the important features belong and y-axis represents a count of the important descriptors within the said descriptor groups **(b)** Top 10 descriptors of the Cas9 classification.

##### Analysis of the Cas9 top 10 ranking important descriptors

We also analyzed the top 10 descriptors of the Cas9 family (Figure 3b). Like Cas12, Schneider-lag2, and normwaalsvolume.G3 appeared in the top 10 important descriptors of Cas9 family. However, in this family, four out of the top 10 descriptors belong to TPC type namely PWN, HHA, DHI, and PYY. This pattern of predominance of TPC descriptors distinguishes Cas9 proteins from Cas12 where Schneider-lag descriptors predominate. To further assess the role of these tripeptides in the Cas9 family, we carried out a Multiple Sequence Alignment (MSA) of the *Streptococcus pyogenes* Cas9 (SpCas9) sequences obtained from UniProt.

##### Tripeptide feature of Cas9

MSA of all Streptococcus Cas9 sequences obtained from UniProt revealed tripeptide PWN to be conserved across all species of SpCas9 proteins. Likewise, the tripeptide HHA is conserved at residues 982-984 in all SpCas9 proteins constituting RuvC region of the sequence. Similarly, DHI and PYY are seen conserved across all SpCas9 proteins forming the HNH and RECIII regions of the sequences. Further analysis revealed that these tripeptides are conserved in other bacterial Cas9 proteins as well. Table 5 summarizes a few bacterial Cas9 species in which we observed the four important tripeptides.

**Table 5:** Examples of Cas9 species containing the four identified tripeptides.

| Residues | Organism | Residue numbers | Domain | PDB |
| --- | --- | --- | --- | --- |
| PWN | SpCas9 | 475-477 | RECIII | 5F9R |
| HHA | SpCas9 | 982-984 | RuvC | 5F9R |
|  | *Neisseria meningitidis*1 Cas9 | 722-724 |  | 6JDV |
|  | *Neisseria meningitidis*2 Cas9 | 722-724 |  | 6JFU |
|  | *Campylobacter jejuni* Cas9 | 706-708 |  | 5X2H |
| DHI | SpCas9 | 839-841 | HNH | 5F9R |
|  | *Actinomyces naeslundii* Cas9 | 581-583 |  | 4OGC |
|  | *Staphylococcus aureus* Cas9 | 556-558 |  | 5AXW |
|  | *Streptococcus thermophilus* Cas9 | 598-600 |  | 6RJD |
| PYY | SpCas9 | 449-451 | RECIII | 5F9R |

Further, we analyzed the distribution patterns of these four tripeptides within our Cas9 training dataset by comparing their descriptor values. As in Figure 4, we observed that all the four TPC descriptors can clearly differentiate between the Cas9 and Non-Cas proteins of our training dataset. It is also seen that the descriptor values of these tripeptides in Cas9 fraction of the training dataset is much higher than their Non-Cas9 fraction thus differentiating the Cas9 proteins from Non-Cas proteins effectively.

**Figure 4.**
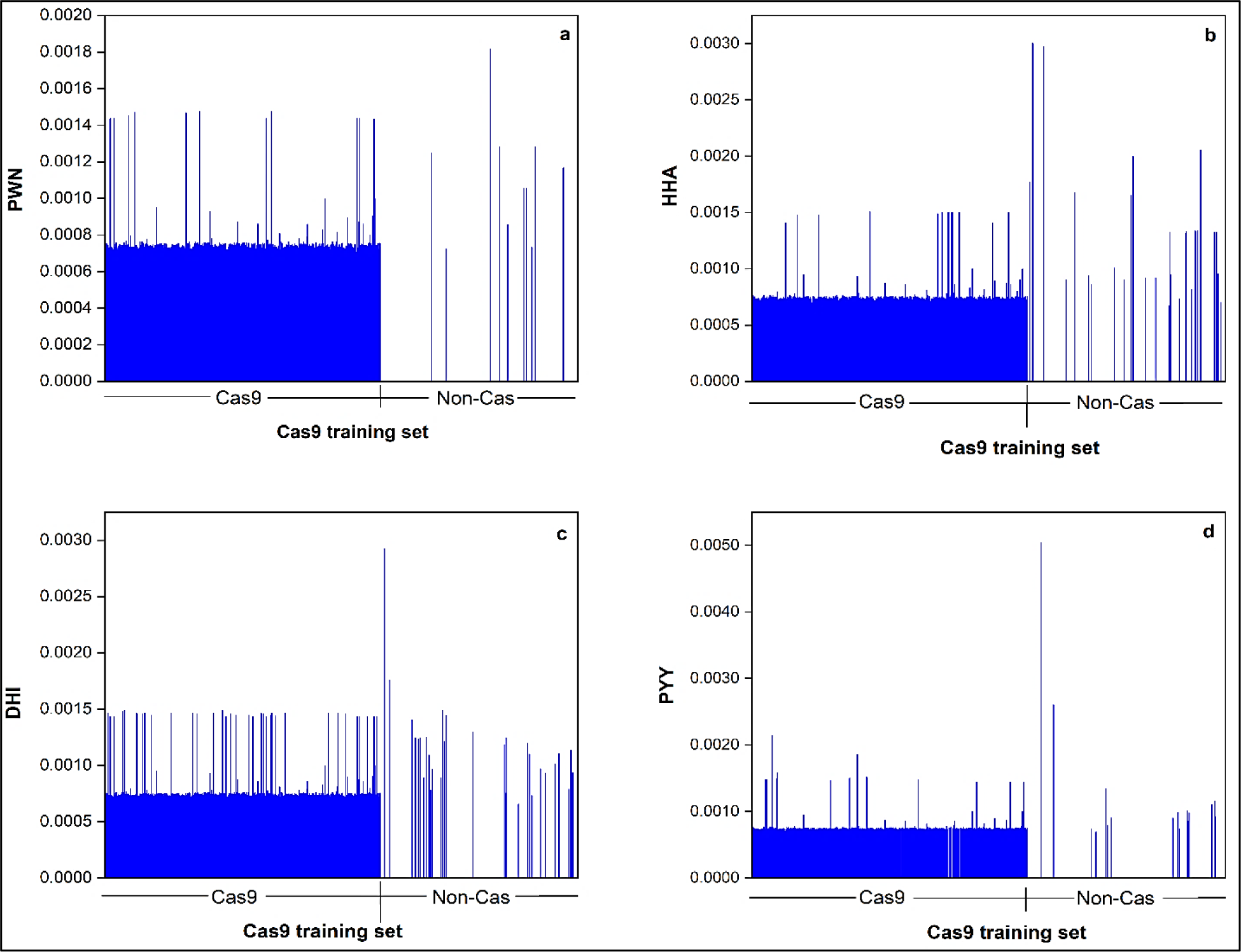
Plot of descriptor values of the four important TPC descriptors **(a)** PWN **(b)** HHA **(c)** DHI **(d)** PYY within the Cas and Non-Cas fractions of our Cas9 training dataset.

Further, we analyzed the percentage distribution of the four tripeptides within the Cas9 and Cas12 training datasets to understand their prevalence patterns in the two Cas families (Figure 5). We observed that the four identified tripeptides (PWN, HHA, PYY, and DHI) distinctly separate Cas12 from Cas9 family. As seen in Figure 5, these peptides are conserved in more than 95% of the Cas9 proteins of the training dataset. This further proves the predominance of tripeptides within the Cas9 family and can thus be useful in distinguishing Cas9 proteins from the other Cas families.

**Figure 5.**
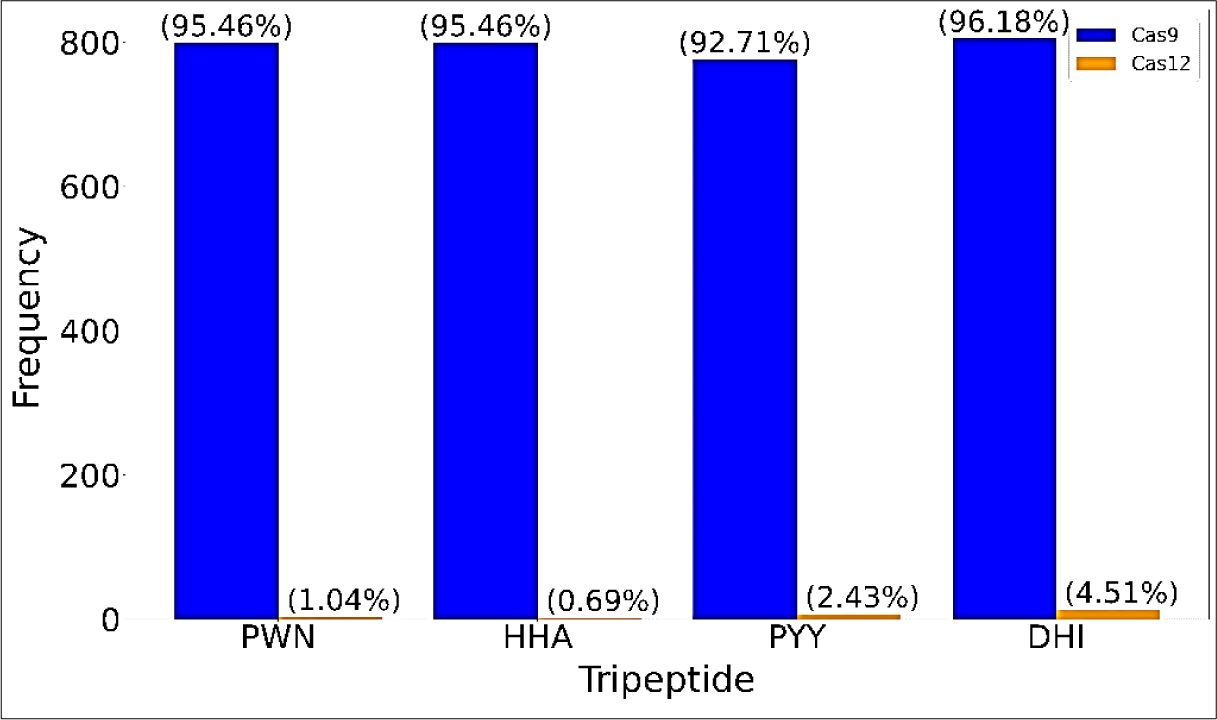
Comparison of prevalence of the four tripeptides identified in Cas9 important features across the two Cas families.

We further tried to investigate the role of these four tripeptides in the SpCas9 crystal structure. Since SpCas9 is the most widely studied Cas9 ortholog for gene editing applications in various living cells and organisms known so far^60–64^, we explored the local arrangements of these four peptides in SpCas9 structure which is shown in Figure 6.

**Figure 6.**
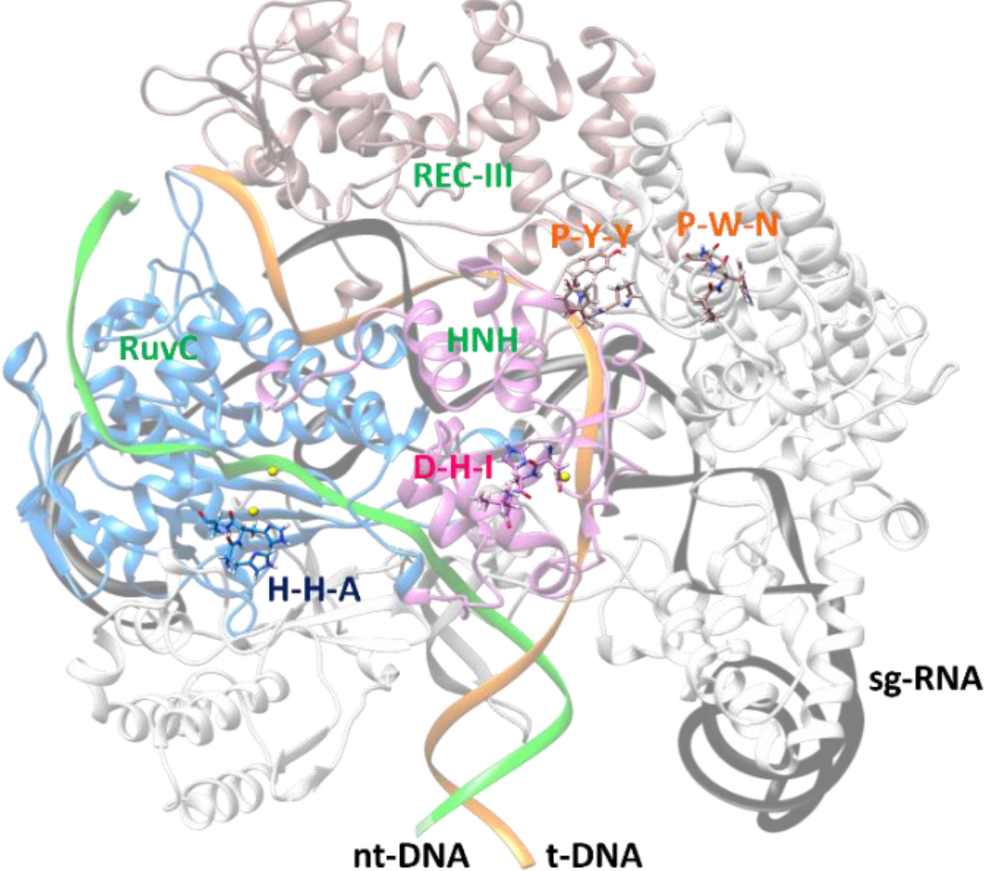
Locations of the four identified tripeptides within the different domains of Cas9 crystal structure [PDB ID: 5F9R]. The tripeptides PWN (475-477) and PYY (449-451) are located in RECIII domain (beigh), while HHA (982-984) and DHI (839-841) are located in RuvC (blue) and HNH (pink) domains of the Cas9 crystal structure.

The tripeptide PWN, and PYY, are located on the REC-III region (of the recognition lobe) of SpCas9 while DHI and HHA are on the nuclease lobe (HNH and RuvC-III). Interestingly, DHI contains two (D838 and H840) of the three catalytic residues required for the target DNA cleavage function of the HNH endonuclease domain of SpCas9^65^. Furthermore, tripeptide HHA (H982,

H983, and A984) is part of the second coordination sphere of the RuvC catalytic site, which facilitates the stability of the RuvC active site for its DNA cleavage function (non-target DNA cleavage). Especially, H939 has been proposed as the catalytic base that is involved in the DNA cleavage mechanism of the RuvC domain^66^. These details ascertain the performance of our ML pipeline and confirm that the identified tripeptides and other top descriptors can add biological insights into the Cas9 catalytic mechanism.

### 3.4. Cas12-Cas9-Non-Cas important features

We further proceeded to identify descriptors that could differentiate between Cas9, Cas12 and Non-Cas proteins. This is a multiclass problem where RF models were developed to classify if a protein sequence is Cas12, Cas9, or a Non-Cas protein. BorutaSHAP selected 118 important features for this dataset which are shown in Figure 7(a). As seen, the top three descriptor sets for the multiclass classification belong to the QSO followed by the C/T/D and then the Conjoint Triad groups. The next set is the SOCNumber which is yet again QSO category of features drawing similarities from the Cas12 and the Cas9 important feature categories.

**Figure 7.**
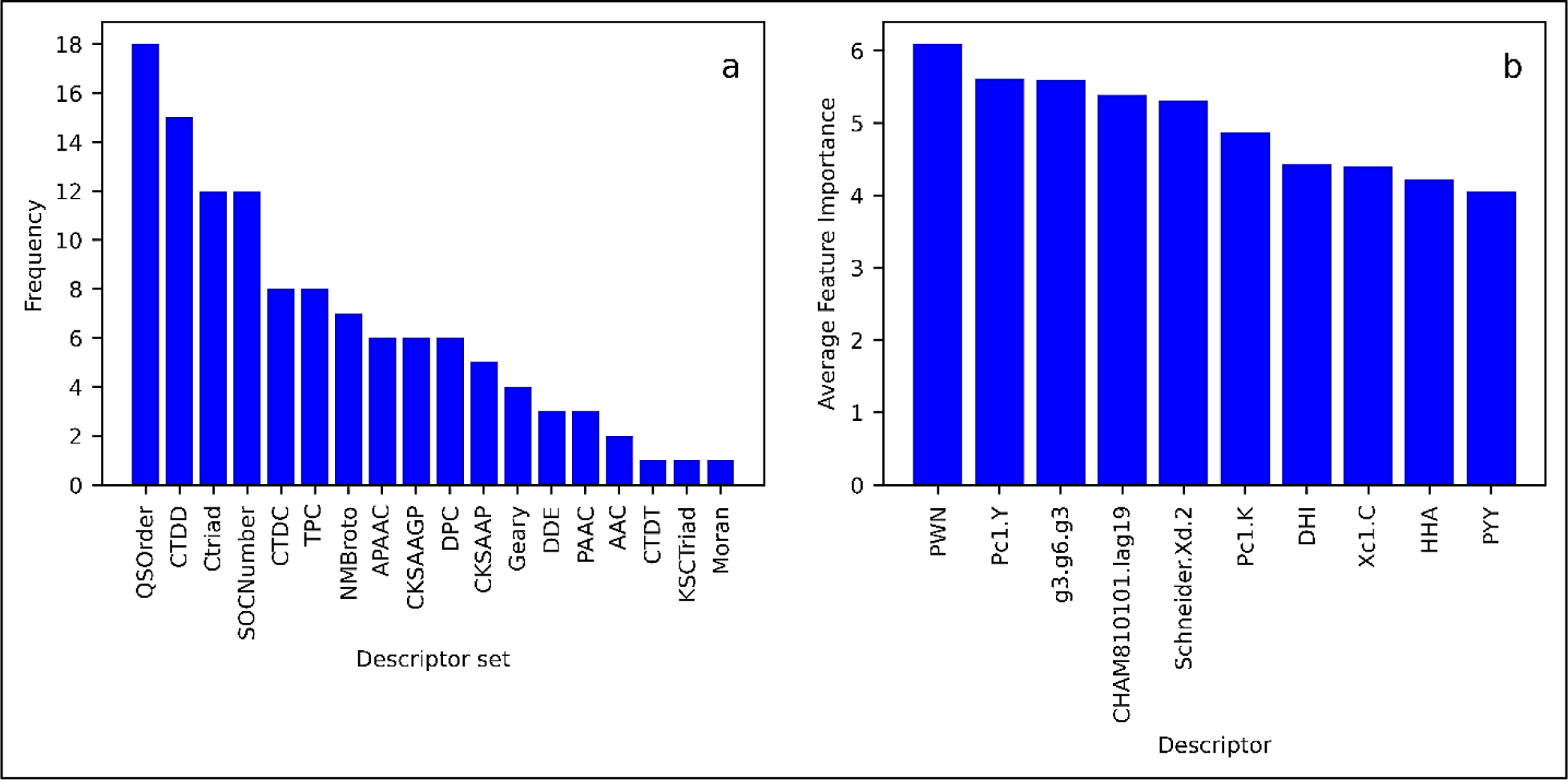
Summary of the important features identified in Cas9 vs. Cas12 vs. Non-Cas multiclass classification. **(a)** Descriptor group-wise distribution of important features identified in multiclass RF model. X-axis represents the descriptor groups to which the important features belong and y-axis represents count of the important descriptors within the said descriptor groups **(b)** Top 10 descriptors of the Cas9 classification.

#### 3.4.1. Analysis of the Cas12-Cas9-Non-Cas classification important descriptors

A deeper look at the top 10 ranking descriptors of the Cas12-Cas9-Non-Cas multiclass classification showed common top features with the previous two binary classification models (Figure 7(b)). The top feature of multiclass classification is “PWN” with the highest feature importance which is also the top feature of the Cas9 family, The next top descriptor is ‘g3.g6.g3’ which is a Conjoint Triad feature (CTF), which was not seen in either Cas12 or Cas9 top descriptors. CTF considers the properties of one residue and its vicinal amino acids and regards three continuous amino acids as a unit^67,68^. Accordingly, the 20 amino acids are categorized into seven groups based upon their physicochemical properties and thereafter, all residues of the same category are considered identical and annotated identically. Thereafter, all sets of three successive amino acids (triad) of a given protein sequence are counted and the triad frequencies are computed over the entire length of the sequence. Based on this annotation, the complete sequence of a protein is encoded as a vector. This method has been successfully utilized in the field of protein-protein interaction prediction^69^. These descriptor values when plotted across the Cas12-Cas9-Non-Cas training dataset, showed an interesting pattern of significantly lower (negative) descriptor values for Cas9 proteins compared to Cas12 and Non-Cas proteins which had positive values for this feature (Figure 8).

**Figure 8.**
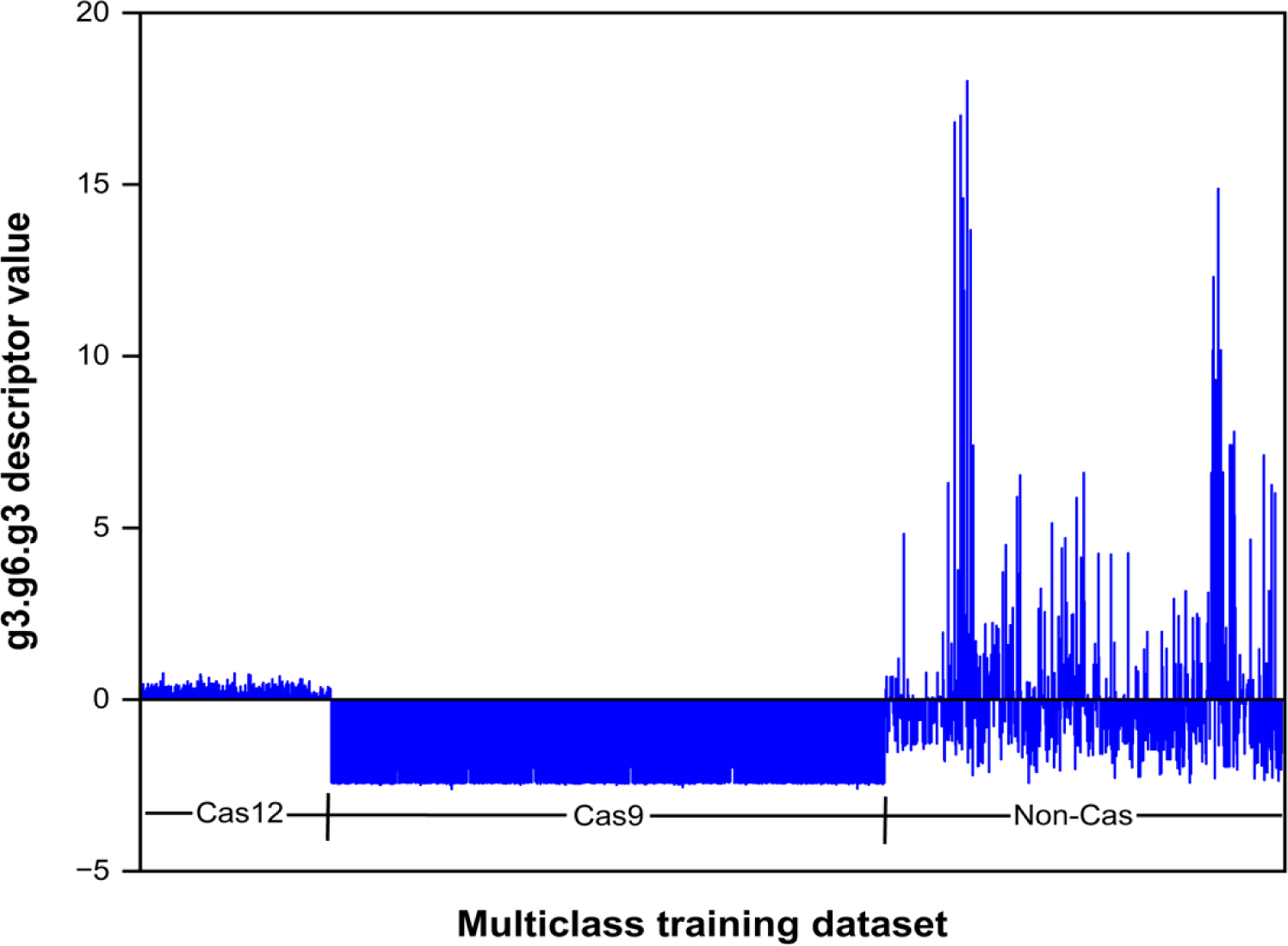
Distribution of the g3.g6.g3 descriptor in the multiclass training dataset

Therefore, this feature can clearly distinguish between the three classes of proteins. This also corresponds well with the fact that the Cas9 or Cas12 top descriptors do not have any Conjoint Triad features making this specific for the multiclass models.

The top 10 important descriptors also include descriptors Pc1.Y and Pc1.K, which are Amphiphilic PAAC (APAAC) descriptors that were first proposed by Chou in 2005 and have since been applied widely to encode different biological sequences like protein and nucleotide sequences. Unlike the traditional AAC features, APAAC descriptors have two components effectively incorporating sequence order effect along with AAC all along the protein sequence. A protein sequence can be represented in terms of APAAC features using the first component which encodes the classic AAC of the amino acids along the protein sequence while the second component incorporates the sequence-order effect using hydrophobicity and hydrophilicity of the amino acids present along the sequence. Given the important roles played by lysine in Cas12 cleavage activity and tyrosine in Cas9 cleavage, these features further emphasize the significance of sequence order information in differentiating the Cas families from Non-Cas proteins. However, the exact role of these two descriptors in the catalytic mechanisms of Cas12 and Cas9 proteins is not yet clearly known. Further, at a global level, we also tried to investigate the contributions of the seven descriptor groups in the three classification tasks. For this, we analyzed the distribution of important features of three classification models across the seven descriptor groups depicted in Table 2.

As seen in Figure 9, the AAC category of descriptors contribute maximum towards the Cas9 important features while QSO group contributes the most for the Cas12 family. For example, seven AAC important descriptors discriminate Cas12 vs Non-Cas while 42 of them discriminate Cas9 vs Non-Cas. Likewise, 31 QSO important features can differentiate Cas12 vs Non-Cas while 19 of them discriminate Cas9 vs Non-Cas. This trend is in accordance with the predominance of Schneider-lag features (QSO) in the Cas12 family and the tripeptide features (AAC) in the Cas9 family seen in the above sections. On the other hand, it is interesting to see that Conjoint Triad descriptors have a large contribution in multiclass but have little to no contributions in Cas12 and Cas9 binary classifications which shows that this feature can clearly differentiate the three families.

**Figure 9.**
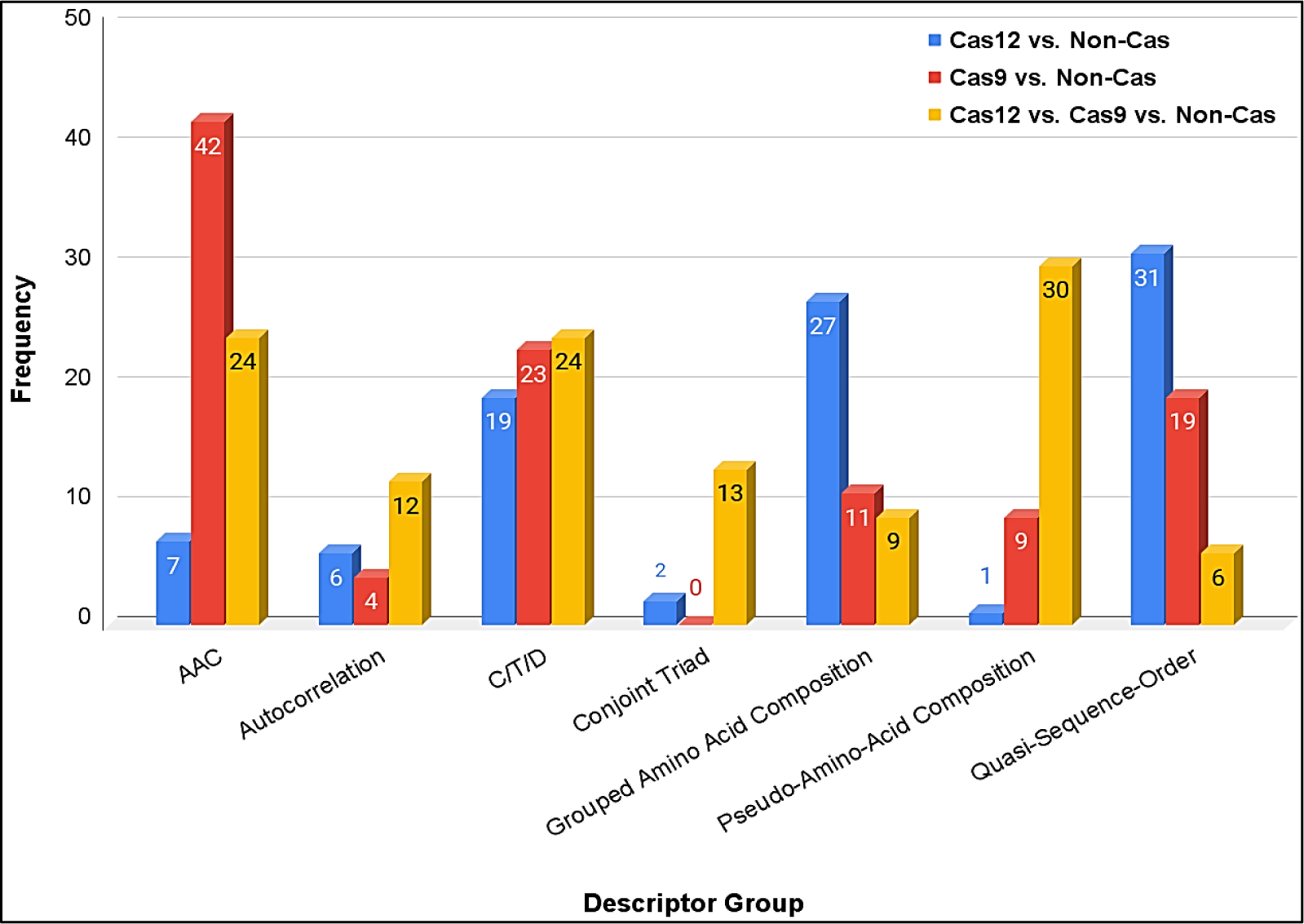
Group-wise distribution of the important features of the three classification models.

##### Comparison with existing Cas prediction tools

There are very few computational tools in literature specifically dedicated to identifying and characterizing Cas proteins. We used the existing tools to compare the performance of our method. We compared our method with CASPredict which is the best-performing existing tool so far in terms of its overall accuracy for Cas protein identification. For comparison, we predicted the independent set of our multiclass dataset on the CasPredict tool using their portal. The comparative performance of our tool with CASPredict is shown in Table 6.

**Table 6:** Comparison of the performance of our method with existing tool.

|  | Accuracy | Specificity | Sensitivity | Precision | F1 score |
| --- | --- | --- | --- | --- | --- |
| CasPredict | 0.861 | 0.878 | 0.853 | 0.941 | 0.895 |
| Our method | **0.987** | **0.993** | **0.989** | **0.986** | **0.987** |

Clearly, our RF models built using the entire protein feature spectrum has outperformed CASPredict. This demonstrated that diverse physiochemical and evolutionary descriptors can contribute significantly to identifying differential patterns and thus aid in Cas protein classification.

## 4. Discussion

In the current study, we developed RF based models that can identify specific Cas9 and Cas12 family of proteins and differentiate them from the Non-Cas proteins. Here Cas9 vs. Non-Cas and the Cas12 vs. Non-Cas classification problems are treated as binary classification problem while the Cas12-Cas9-Non-Cas classification is treated as a multiclass problem. The RF classifiers are built using a multitude of protein features which encode for the physiochemical, evolutionary, and compositional information of the proteins and hence hold the ability to accurately capture detailed patterns in the Cas protein families. Feature selection for both the binary and the multiclass classification is done using the BorutaSHAP feature importance method with a 5-fold data split incorporated during feature selection. Our models were evaluated rigorously on test set followed by prediction on independent set across 15 different data splits using six different evaluation metrices. On the test set, the binary models of Cas12 vs Non-Cas and Cas9 vs Non-Cas achieved a high accuracy close to 96% and 99%. The multiclass model also showed a high F1 score of close to 99%. The models performed equally well on the independent dataset achieving a high accuracy of around 92% and 95% and an F1 scores of around 97% for the multiclass models. These results show the high performance of our models. Our models developed using the diverse feature set, clearly outperformed existing Cas prediction tool like CasPredict which was built using only DPC based features. Through this study, we also understand the descriptors specific to the two Cas families which can assist in the identification and differentiation of Cas12 and Cas9 proteins.

Analysis of important descriptors of both the binary classifications showed that QSO descriptors are important for the Cas12 family while AAC features are important for Cas9 family. Among the AAC group, TPC descriptors emerged to be highly important, which are routinely used in various protein classification and functional characterization tasks. One of the most important features in Cas12 family is “Polarizability.G3” which indicates that the global composition of group 3 polarizability residues in Cas12 proteins could be beneficial to the Cas12 family. Sequence order information features like SQCNumber or QSOrder emerged to be highly important for Cas12 families. Especially the “Schneider-lag” based features which depict the coupling modes between contiguous residues separated by certain distance along the length of protein sequence emerged to be predominantly important for Cas12 family. These features capture relationships between residue pairs located at various positions along the length of a protein sequence and may therefore explain their long-range interactions which provide insights about the coevolved residues driving protein-protein interactions. In addition to the sequence order features, Cas12 family proteins are also dominated by composition features like charge, volume and polarizability. In the Cas9 family, four specific tripeptides “PWN”, “HHA”, “PYY” and “DHI” emerged to be very important. MSA of Cas9 sequences along with the analysis of literature and existing Cas9 structures revealed these tripeptides to constitute catalytically important domains of Cas9 strucutres. The four tripeptides could clearly differentiate between Cas9 and Cas12 families with a significantly higher content of these tripeptides in Cas9 than Cas12 proteins. Further, in SpCas9 crystal structure, PWN and PYY formed the recognition lobe which is important in binding the sgRNA and DNA. More remarkably, one of our identified tripeptides DHI contains two (D838 and H840) of the three catalytic residues required for target DNA cleavage function of HNH domain of SpCas9.

Overall, sequence order features like Schneider-lag features accounting for specific short- and long-range interactions and QSO features along with CTDC features like charge, normalized Van der Walls volume and polarizability characterized Cas12 family. On the other hand, Amino acid composition (especially TPC) and PAAC descriptors can explain the Cas9 family well. The four tripeptides lie in various catalytically important domains of SpCas9 and aid in binding and cleavage activities. These are also predominantly distributed and conserved across the Cas9 proteins. This is the first study demonstrating the successful application of vivid protein features to classify Cas proteins and conduct an in-dept investigation of the protein features governing the Cas9 and Cas12 families. We strongly believe that our analyzed features can guide future structural biologists and computational biologists to design novel Cas proteins with desired properties like optimized size, efficient delivery of CRISPR machinery, improved cleavage efficiency, and reduced off-target effects.

## 5. Conclusions

This study demonstrates a systematic methodology of Cas protein classification using a multitude of protein features to identify descriptors which are specific to each Cas families. Here we demonstrate our method on two Cas families; Cas9 and Cas12. A multiclass classification is also built to ensure that the classifier could clearly differentiate between the Cas9, Cas12, and Non-Cas families. The developed models were evaluated on the test set followed by independent set prediction. To account for the variance in our models, we rigorously repeated feature selection, training, tuning, and testing over 15 different data splits. All three classification models performed extremely well on the test set as well as the independent sets achieving accuracies close to 96%. A deeper investigation of the top 10 features highlighted the protein features specific to the Cas9 and Cas12 families. For the Cas12 family, QSO descriptors, especially the Schneider-lag descriptors which can explain the long- and short-range interactions in proteins emerged important. Along with QSO, Composition-based features like charge, polarizability, and Van der Walls volume are also significant. On the other hand, amino acid composition viz. tripeptide features dominated the Cas9 family. Specifically, four tripeptide descriptors PWN, HHA, DHI, and PYY emerged to be extremely important. Further analysis showed that these four tripeptides are conserved across all the Cas9 species. Moreover, these tripeptides are seen to be located within the catalytically important domains of SpCas9 crystal structure demonstrating the importance of the identified tripeptides. Interestingly, two of the tripeptides HHA and DHI are identified to be actively involved in Cas9 DNA binding and cleavage activities. In line with that, we propose PYY and PWN, two of the 4 identified tripeptides located in the RecIII region to be extremely important for the Cas9 family and may perhaps be involved in the recognition activity of the RECIII domain. Therefore, our proposed pipeline can not only identify and characterize novel Cas9 and Cas12 proteins but also guide further modifications of Cas proteins to optimize their cleavage activities.

## Acknowledgments

This work was supported by a grant from National Institute of General Medical Sciences of the National Institutes of Health (R35GM133657). We thank our collaborator professor Shouyi Wang and his group from the University of Texas Arlington (UTA) for their continuous and valuable research support all along this study.

## Data Availability

The training and independent protein sequences for all the three datasets Cas9, Cas12 and Non-Cas are given as supporting information. The global ranking for all the intersection features is all the three classifications is also given as supporting information.

## Author Information

### Authors

**Sita Sirisha Madugula**-*Department of Pharmaceutical Sciences, University of North Texas System College of Pharmacy, University of North Texas Health Science Center, Fort Worth, Texas, 76107, United States*

**Pranav Pujar-***Department of Industrial, Manufacturing and Systems Engineering, The University of Texas at Arlington, 701 South Nedderman Drive, Arlington, Texas, 76019*

**Nammi Bharani**-*Department of Industrial, Manufacturing and Systems Engineering, The University of Texas at Arlington, 701 South Nedderman Drive, Arlington, Texas, 76019*

**Shouyi Wang-** Department of Industrial, Manufacturing and Systems Engineering, University of Texas at Arlington, 701 South Nedderman Drive, Arlington, Texas, 76019, United States.

**Vindi Mahesha Jayasinghe Arachchige**-*Department of Pharmaceutical Sciences, University of North Texas System College of Pharmacy, University of North Texas Health Science Center, Fort Worth, Texas, 76107, United States*

**Tyler Pham-***Graduate School of Biomedical Sciences, University of North Texas Health Science Center, Fort Worth, Texas, 76107, United States*

**Dominic Mashburn-** *Department of Pharmaceutical Sciences, University of North Texas System College of Pharmacy, University of North Texas Health Science Center, Fort Worth, Texas, 76107, United States,*

**Maria Artilis-** *Department of Pharmaceutical Sciences, University of North Texas System College of Pharmacy, University of North Texas Health Science Center, Fort Worth, Texas, 76107, United States,*

## Author Contributions

JL and SW conceived the idea, provided the supported to develop the workflow, reviewed the manuscript critically and monitored over the study. MSS designed the study, developed the ML models, contributed to result analysis, and drafted the manuscript in coordination with other authors. PP contributed by calculating all the protein features required for developing ML models. NB contributed by giving research ideas during ML model development. VMJA and TP collected and cleansed the Cas9, Cas12 and Non-Cas datasets. DM and MA assisted and provided research inputs in result analysis and writing the discussion section. They helped in literature survey to understand the relevance of the obtained important descriptors for each CAS family. All the authors have reviewed the manuscript critically and given intellectual input collectively contributing to the study.

## Notes

The authors declare no competing financial interests.

