## Supplementary information for "Identification of Family-Specific Features in Cas9 and Cas12 Proteins: A Machine Learning Approach Using Complete Protein Feature Spectrum"

| Table of Contents | Page No. |
| --- | --- |
| **Table S1.** Summary of the different hyperparameters and their ranges tuned in the study | S3 |
| **Table S2.** Performance of the three classification models on the test set using union features averaged over 15 runs. | S4 |
| **Table S3.** Performance of the three classification models on the independent set using union features averaged over 15 runs | S5 |

**Table S1.** Different hyperparameters and their ranges tuned in the study.

| **Parameter** | **Range** |
|---|---|
| n_estimators | 10,20,30,40,50, 60,70, 80,90,100,110, 120,130,140, 150 |
| max_depth | 10,20,30,40,50, 60,70, 80,90, 100 |

**Table S2:** Performance of the three classification models on the test set using union features averaged over 15 runs.

| | **Accuracy** | **Precision** | **Recall** | **F1-score** | **AUC** | **Specificity** |
|---|---|---|---|---|---|---|
| Cas12 vs. Non-Cas | 0.96741± 0.014 | 0.9686± 0.015 | 0.9574± 0.020 | 0.9623± 0.017 | 0.9574± 0.020 | 0.9861± 0.012 |
| Cas9 vs. Non-Cas | 0.9974± 0.002 | 0.9976± 0.002 | 0.9971± 0.003 | 0.9973± 0.002 | 0.9971± 0.003 | 0.9950± 0.005 |
| Cas12-Cas9-Non-Cas | 0.9982± 0.002 | 0.9986± 0.001 | 0.9985± 0.002 | 0.9985± 0.002 | 0.9996± 0.0008 | 0.9989± 0.001 |

**Table S3:** Performance of the three classification models on the independent set using union features averaged over 15 runs.

| | **Accuracy** | **Precision** | **Recall** | **F1-score** | **AUC** | **Specificity** |
|---|---|---|---|---|---|---|
| Cas12 vs. Non-Cas | 0.9177± 0.081 | 0.9407± 0.050 | 0.9098± 0.089 | 0.9112± 0.092 | 0.9098± 0.089 | 0.9986± 0.005 |
| Cas9 vs. Non-Cas | 0.9575± 0.027 | 0.9584± 0.024 | 0.9570± 0.030 | 0.9570± 0.028 | 0.9570± 0.030 | 0.9625± 0.0208 |
| Cas12-Cas9-Non-Cas | 0.9746± 0.018 | 0.9754± 0.016 | 0.9751± 0.019 | 0.9749± 0.001 | 0.9962± 0.001 | 0.992± 0.01 |
